# Co-Activation Patterns Characterize Early Resting-State Networks in Newborn Infants: A High-Density Diffuse Optical Tomography Study

**DOI:** 10.1101/2025.03.15.643385

**Authors:** K. Lee, J. Uchitel, C. Caballero-Gaudes, L. Collins-Jones, R. Cooper, A. Edwards, J. Hebden, G. Kromm, K. Pammenter, T. Austin, B. Blanco

**Author notes:** Corresponding Author: Borja Blanco.

## Abstract

**Significance:** Dynamic functional connectivity (FC) in neonates is a growing area of interest due to the developmental significance of early functional networks. There are several emerging techniques to measure dynamic FC, adding new perspectives to well-studied static FC networks. Recent dynamic FC studies suggest that adult resting-state networks are driven by key moments of dynamic activity rather than sustained correlations. Co-activation pattern (CAP) analysis leverages this theory, clustering high-activity frames to identify recurring configurations of significant activity. High-density diffuse optical tomography (HD-DOT) is an infant-friendly modality that measures hemodynamic changes in the cortex and has been used to investigate static FC in term-aged infants. CAP analysis has not yet been applied to neonatal HD-DOT and may reveal temporal features of early functional networks that are not apparent from static methods.

**Aim:** This study applies CAP analysis to neonatal HD-DOT to characterize transient co-activation states and provide new insight into early functional brain networks beyond what is known from static FC methods alone.

**Approach:** Task-free HD-DOT data were acquired from a cohort of sleeping term newborns at the Rosie Hospital, Cambridge UK (n = 44, postmenstrual age = 40+3 (range: 38+2–42+6) weeks). In each recording, the top 15% of seed-selected frames were clustered using the K-means algorithm for three regions of interest (ROIs: frontal, central, and parietal) to identify significant seed-associated patterns of co-activation or co-deactivation. These co-activation patterns (CAPs) were characterized for each infant using four metrics: consistency, fractional occupancy, dwell time, and transition likelihood.

**Results:** Distinct CAPs, reflecting the dynamic organization of neonatal cortical networks, were identified for frontal, central, and parietal regions. These CAPs showed high consistency scores, reflecting high intra-cluster spatial correlation and validating the efficacy of CAP analysis for newborn HD-DOT data. The CAP decomposition revealed significant patterns not observed in conventional static functional connectivity analyses. Several neonatal CAPs exhibited frontoparietal co-activation, potentially reflecting early default mode network activity, which is immature and modular for the first year after birth.

**Conclusions:** This work demonstrates the utility of CAP analysis with newborn HD-DOT and provides new insight into the dynamics of neonatal functional connectivity.

## 1. Introduction and Background

The first year after birth is marked by significant structural and functional changes in the human brain (Keunen et al., 2017; Stoecklein et al., 2020). Studying functional maturation in the neonatal period is critical for characterizing patterns of typical and atypical brain development and their association with lifelong neurodevelopmental variation. Despite the importance of understanding early brain development, neonatal studies remain far less common than adult studies, restricting insight into how functional brain networks emerge and mature during this critical period and limiting our ability to translate advanced analytical methods to infant populations. This disparity likely stems from the challenges of imaging the newborn brain, including practical, technological and analytical considerations unique to neonatal participants (Mohammadi-Nejad et al., 2018).

Although functional magnetic resonance imaging (fMRI) has greatly advanced our knowledge of early brain development and is generally regarded as the gold standard for functional neonatal brain imaging, fMRI has important limitations (Edwards et al., 2022; Eyre et al., 2021; Keunen et al., 2017). fMRI requires motion-free scans, and neonates (asleep or awake) move frequently and unpredictably, restricting prolonged or repeated scanning (Arthurs et al., 2012; Mohammadi-Nejad et al., 2018). It is for this reason that neonates used to be sedated before fMRI scans, which was suboptimal as sedation has been found to modulate brain activity (Gemma et al., 2009; Hassanzadeh et al., 2023; Keunen et al., 2017; Mohammadi-Nejad et al., 2018). Despite advances in protocols for scanning awake infants (Deen et al., 2017), fMRI is not conducive to naturalistic studies as neonates are scanned in a loud machine isolated from their mothers, disrupting their natural environment (Arthurs et al., 2012; Mohammadi-Nejad et al., 2018). Wearable high-density diffuse optical tomography (HD-DOT) systems have been recently developed to facilitate infant studies because of their comfortable design, relatively increased resilience to motion artifacts, and cot-side applicability (Collins-Jones et al., 2024; Frijia et al., 2024; Uchitel et al., 2023). HD-DOT is an extension of functional near-infrared spectroscopy (fNIRS) that uses dense arrays of overlapping, multi-distance sources and detectors to reconstruct three-dimensional maps of cortical oxygenated ([HbO]) and deoxygenated ([HbR]) hemoglobin concentrations. HD-DOT can image changes in [HbO] and [HbR] with higher temporal resolution than fMRI and has a superior spatial resolution compared to electroencephalography (EEG), though, like EEG, it is limited to cortical imaging (Peng & Hou, 2021; Uchitel et al., 2023). These features of wearable HD-DOT have advanced the study of newborn infant functional connectivity across a range of contexts, including clinical and home settings (Uchitel et al., 2023, Clackson et al., 2025).

Functional connectivity is a central concept in the study of typical and atypical brain development, describing the functional relationship between spatially distinct brain regions. It is commonly defined as the strength of the covariance or correlation of activity between brain regions over time (Power et al., 2010). Functionally connected brain regions may be spatially distant yet exhibit temporally coupled activity (Arimitsu et al., 2022). Functional brain networks are groups of interconnected regions and are associated with specific neurophysiological processes (Menon, 2013; Power et al., 2010). fMRI has reliably identified several resting-state networks in the adult brain (e.g., default-mode network, primary sensorimotor network, visual network, and language network) (M. H. Lee et al., 2013; J. Yang et al., 2020). Although much remains unknown about the emergence of these networks, several fMRI studies have explored functional connectivity in the infant brain. Primary sensorimotor and visual networks are present at birth, and additional networks gradually emerge during the first year, with primary networks developing earlier than higher-order networks (Gao et al., 2017). By age one, several resting-state networks resemble their analogous adult networks, including primary and secondary visual networks, dorsal attention, and default mode network. Still, some networks are underdeveloped by age one, such as the frontoparietal executive control network (Gao et al., 2017).

These studies provide important insight into early brain development, however, they approach functional connectivity from the conventional “static” point of view. Static functional connectivity refers to the average functional connectivity over the course of the data collection window. This traditional approach is robust but does not capture the dynamic, time-varying nature of functional connectivity, which is inherently non-stationary. Dynamic functional connectivity studies aim to describe time-varying functional interactions between brain regions, adding depth and accuracy to our understanding of functional brain organization (Hutchison et al., 2013; Preti et al., 2017). In recent years, many fMRI, EEG, and fNIRS studies have explored methods to measure dynamic functional connectivity. Several groups have explored different measures of dynamic functional connectivity such as neural flexibility, temporal variability, and stability of correlation coefficients (Hutchison et al., 2013; Preti et al., 2017; Wen et al., 2020; Yin et al., 2020). Notably, these dynamic features of functional connectivity change with age (Qin et al., 2015) and can be used to differentiate adult controls from patients diagnosed with schizophrenia (Shen et al., 2014) or Alzheimer’s disease (Jones et al., 2012).

In contrast to static functional connectivity measures like seed-based correlation or independent component analysis (ICA), which require clean and continuous data segments, dynamic functional connectivity analysis operates on a finer time scale and is therefore more tolerant of fragmented datasets. This flexibility makes it a promising approach for neonatal studies, which often encounter motion artifacts or are limited to short scanning sessions. Several groups have investigated dynamic functional connectivity in neonates, characterizing resting-state networks in term and preterm infants. Their methods vary, drawing upon analogous adult studies in the fMRI community. Three neonatal studies calculated dynamic functional connectivity using a sliding-window approach, temporally partitioning the BOLD signal into overlapping or non-overlapping segments and then calculating parcel or voxel correlation matrices in these windows (Niu et al., 2020; Ren et al., 2022; Wang et al., 2024). Another common method applies clustering algorithms, such as K-means, to windowed ICA time courses to identify recurring patterns of network interactions (López-Vicente et al., 2021; Ma et al., 2020; Marusak et al., 2017).

These frameworks identify recurring patterns of brain activation and characterize their variation over time. As with many window-based analyses, this method is sensitive to changes in analysis parameters such as window and overlap length which cannot be empirically chosen (Cabral et al., 2017; Hutchison et al., 2013). To address the issue of parameter sensitivity, a new analysis methodology was developed for fMRI data that does not depend on time-domain parameters (Liu and Duyn, 2013). This method revealed that resting-state brain activity is driven by distinct, instantaneous configurations occurring at single time points. These recurring configurations are referred to as co-activation patterns (CAPs) and can be identified using unsupervised clustering techniques. The spatial patterns of CAPs’ describe significant arrangements of instantaneous brain activity, while metrics such as dwell time and fractional occupancy, capture their dynamic properties. CAPs and their associated metrics have been used in adult and pediatric populations to examine task, state, and cohort differences (Adhikari et al., 2021; Marshall et al., 2020; Mawla et al., 2023; H. Yang et al., 2021). However, they have not yet been applied to neonatal populations, leaving a gap in our understanding of early dynamic functional connectivity. The premise of CAP analysis conceptually and procedurally resembles that of EEG microstate studies, which have successfully characterized resting-state brain dynamics in neonates and infants, revealing similarities with adult microstates and providing insight into global network dynamics in preterm neonates (Brown & Gartstein, 2023; Hermans et al., 2023).

CAP analysis in HD-DOT remains limited (Khan et al., 2022), and, to our knowledge, it has not been applied to neonatal HD-DOT data. Addressing this gap, the present study has two objectives: first, to validate CAP analysis for neonatal HD-DOT; and second, to shed light on early dynamic functional connectivity by presenting the first HD-DOT CAP decomposition of task-free networks in term-aged neonates. This approach provides a novel opportunity to capture the transient, time-varying organization of the neonatal brain, which is inaccessible to traditional static analyses.

## 2.0 Methods

### 2.1 Participants

This study was approved by the National Research Ethics Service East of England Committee (REC reference 15/LO/0358). Written informed consent was provided by the parents of 82 healthy newborn infants (born ≥ 37 weeks’ gestation) at Rosie Hospital (Cambridge University Hospitals NHS Foundation Trust). 38 neonates were excluded from the study post-data collection due to insufficient quantity of motion-free data (see Section 2.3), consistent with the exclusion rate typically observed in fNIRS infant studies (C. W. Lee et al., 2017; Uchitel et al., 2023). Data from 44 neonates were preprocessed and analyzed (n female = 15; median gestational age [GA] = 40+0 (range: 38+1–42+1) weeks; median postmenstrual age [PMA] = 40+3 (range: 38+2–42+6) weeks; median age at time of study = 2 (range: 0–11) days; median birth weight = 3.405 (range: 2.415–4.240) kg; median head circumference = 34.5 (range: 33.0–37.0) cm. Twenty-eight of the 44 neonatal datasets were previously used by Uchitel et al., (2023) to examine static functional connectivity.

### 2.2 Data Collection

HD-DOT data were collected cot-side using a modular wearable HD-DOT technology called LUMO (Gowerlabs Ltd, UK, https://www.gowerlabs.co.uk/). Gowerlabs assisted in the construction of LUMO caps customized for two neonatal head circumferences (34 cm and 36 cm) which utilized neonatal EasyCaps (Easycap GmbH, Germany). These caps were fitted with 12 HD-DOT tiles, each containing 3 sources and 4 detectors (36 sources and 48 detectors in total), and were designed to cover frontal, central, and parietal regions of interest (ROIs). The neonatal LUMO caps were capable of collecting intensity data for two wavelengths (735 nm and 850 nm) at a sampling rate of 10 Hz.

Data collection followed the protocol outlined previously (Uchitel et al., 2023). Neonates were recorded while sleeping in their cot on the postnatal ward. Parents were typically present throughout the duration of the study. If the infant was restless, they were swaddled and/or fed to encourage sleep. Although recordings were ideally planned for one hour or longer, some sessions ended early due to infants waking up, wanting to feed, or undergoing clinical procedures. Synchronized video of the infant was captured to confirm the neonate remained asleep during acquisition. Lastly, a 3D scan of the infant’s head was collected using the “Scandy Pro” iPhone application (Scandy LLC, USA) for digitization of head landmarks and tile locations. These locations were used to scale the neonatal head model for image reconstruction.

### 2.3 HD-DOT Data Preprocessing

Data preprocessing for this study was based on the framework used in a previous HD-DOT infant study (Uchitel et al., 2023), with modifications to better suit the current study objectives. All preprocessing was done in MATLAB R2021b (Mathworks, USA) using a mixture of in-house scripts and the following fNIRS MATLAB toolboxes: Homer2 (Huppert et al., 2009) and DOT-HUB (https://github.com/DOT-HUB). Motion-free data segments (duration ≥ 100 seconds) were extracted from each participant’s dataset using the *hmrMotionArtifact* function from Homer2. Participants with less than three minutes total of motion-free data were excluded from further analysis. Channels were evaluated using a sliding window approach (window = 33% the length of the segment, overlap = 50%) and were removed if their coefficient of variation exceeded 12 or if their average signal was outside the expected signal range (1e-4 ≤ mean signal ≤ 2.5). Data were converted from optical density to [HbO] and [HbR] using the *hmrOD2Conc* function from Homer2. Non-neuronal hemodynamic sources of noise such as scalp-hemodynamics were regressed out of the concentration data using short separation regression (Sato et al., 2016), a process which regresses out the average activity of short distance channels activity to attenuate scalp-level physiological confounds. Attenuating physiological confounds increases the likelihood of obtaining measures from cerebral rather than extra-cerebral origin (Yücel et al., 2021). Temporal filtering was performed in a linear regression model. Contributions of high-frequency physiological noise sources such as respiration and cardiac pulsation were accounted for in the model by including Fourier terms for frequencies above 0.08 Hz (Blanco et al., 2021). Slow frequency fluctuations and signal drifts were also accounted for in the model by adding the kth order Legendre polynomials (where k = 1 + floor (duration in seconds / 150)) because their spectral power predominantly consists of very low frequencies. Concentration data were converted back to optical density using the *hmrConc2OD* function from Homer2. Lastly, since hemodynamic changes in the brain happen on the scale of seconds, the data were downsampled to 1 Hz to reduce computational load of data processing.

### 2.4 HbO and HbR Image Reconstruction

The source and detector locations of each tile were marked by fluorescent triangle stickers on the HD-DOT cap. This allowed for manual identification of each source and detector in the photogrammetry Scandy Pro head scans utilizing a software called CloudCompare (https://www.cloudcompare.org/). CloudCompare rigidly transformed the manually identified point clouds to create a complete head model of the sources, detectors, and head landmarks (nasion, inion, Cz, and pre-auricular locations). Next, an affine transformation (computed using head landmarks positions) was used to register the source and detector coordinates to an infant head model (*DOTHUB_LUMO2NIRS* function from the DOTHUB toolbox). This model was drawn from a neonatal head model database and selected to match the average head circumference of the cohort (Collins-Jones et al., 2021).

Images of brain activity were reconstructed using DOTHUB and Toast++ MATLAB toolboxes (Schweiger & Arridge, 2014; Uchitel et al., 2023). The Jacobian matrix (calculated using Toast++ toolbox) was first constructed to be 30 x 30 x 30 voxels and then projected into a 3D infant-specific head mesh. To model [HbO] and [HbR] concentration, tissue absorption at wavelengths 735 and 850 nm were specifically modeled. Changes in absorption coefficients at these wavelengths were calculated by inverting the zeroth-order forward model (calculated using diffusion approximation) (Arridge, 1999; Arridge & Schotland, 2009). These relative absorption coefficient differentials were used to create images of changes in [HbO] and [HbR]. Images were mapped from the volume mesh (626580 nodes) to the surface mesh (34194 nodes) of the head model for cortical visualization.

A group-level sensitivity mask was calculated by finding the sensitivity for each surface node at the participant level using the *DOTHUB_MakeGMSensitivityMap* function. A node was considered sensitive if it exhibited a sensitivity greater than 10% the maximum value of the normalized Jacobian. A final group-level sensitivity mask was created by identifying nodes to which at least 75% of subjects were sensitive (see Figure 2a).

**Figure 2.**
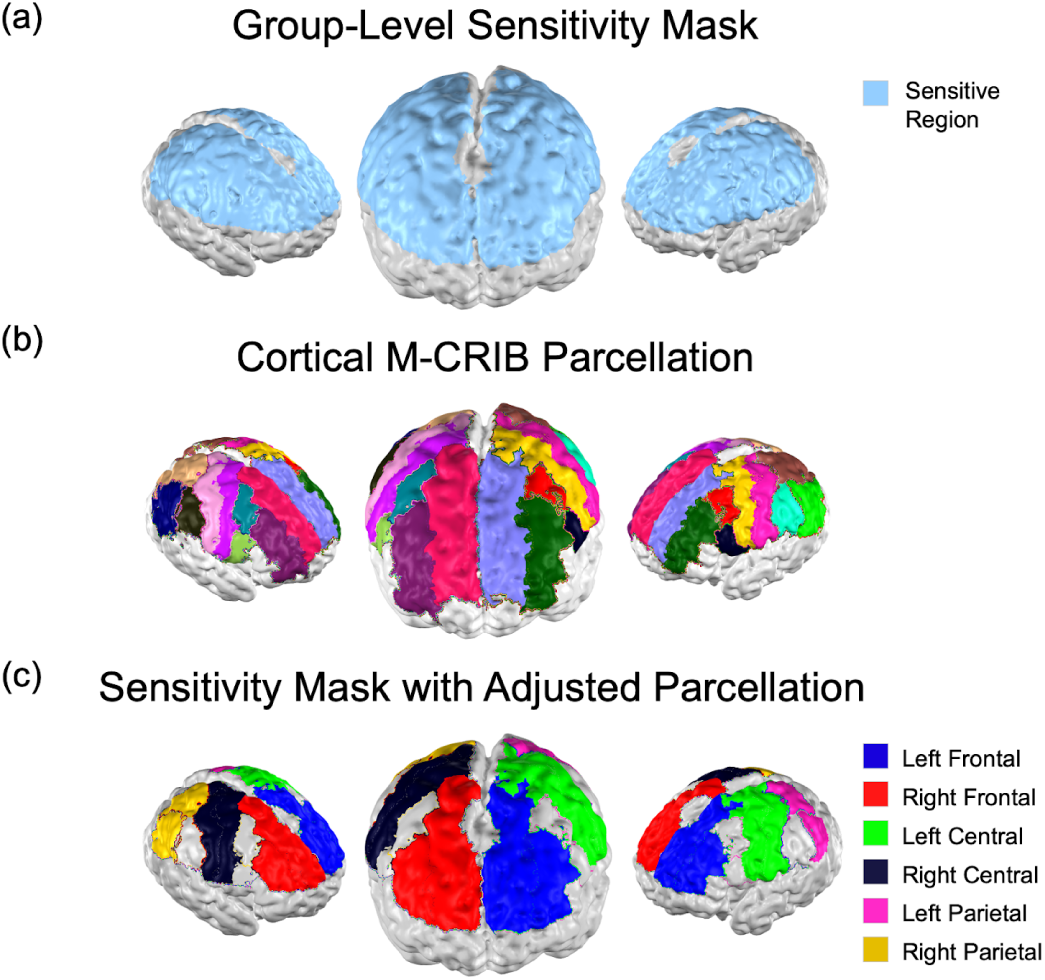
(a) Group-level sensitivity profile co-registered to neonatal head model. (b) M-CRIB parcellation co-registered to the neonatal head model. (c) Group-level sensitivity profile parcellated by M-CRIB labels adjusted for regions of interest.

### 2.5 Parcellation and Seed Selection

This study aimed to examine brain activity in the three cortical regions covered by the HD-DOT system: frontal, parietal, and central regions. These regions were selected to interrogate different and distinct functions (higher executive, somatomotor, and visuospatial processes) which have been well-researched in analogous static functional connectivity studies. Regions were identified using the M-CRIB neonatal parcellation atlas (Alexander et al., 2019, see Figure 2b). The M-CRIB atlas supplied labels for 9 regions per hemisphere. Left and right rostral middle frontal gyrus and superior frontal gyrus labels were combined to create left and right frontal regions respectively. Left and right precentral gyrus and postcentral gyrus labels were combined to create left and right central regions respectively. Left and right superior parietal gyrus and inferior parietal gyrus were combined to create left and right parietal regions respectively. In total, six brain regions were studied: left and right frontal, central, and parietal regions (see Figure 2c). After seed selection, left and right homologous regions were combined and treated as a single ROI.

Since this study relies on seed-based frame selection, it was critical to choose seeds which highlighted activity in the corresponding region. This motivated our decision to calculate optimal seeds at the participant level rather than select a singular seed driven by group-averaged data (see Figure 3b and Supplementary Figure 1). For each brain region, all nodes within that region were considered as candidate seed locations. For each candidate seed, an [HbO] time course was computed by averaging the activity of nodes within a 3 mm radius. This time course was then correlated with the rest of the cortical mesh to generate a correlation map. To determine which candidate best represented the brain region, each correlation map was thresholded (r > 0.5) and compared with a mask of the brain region and its bilateral counterpart using the Dice similarity coefficient. The candidate seed with the highest Dice score, indicating the strongest correspondence with the brain region, was selected as the regional seed for that individual. This procedure was repeated for all six brain regions (seed locations are shown in Supplementary Figure 1).

**Figure 3.**
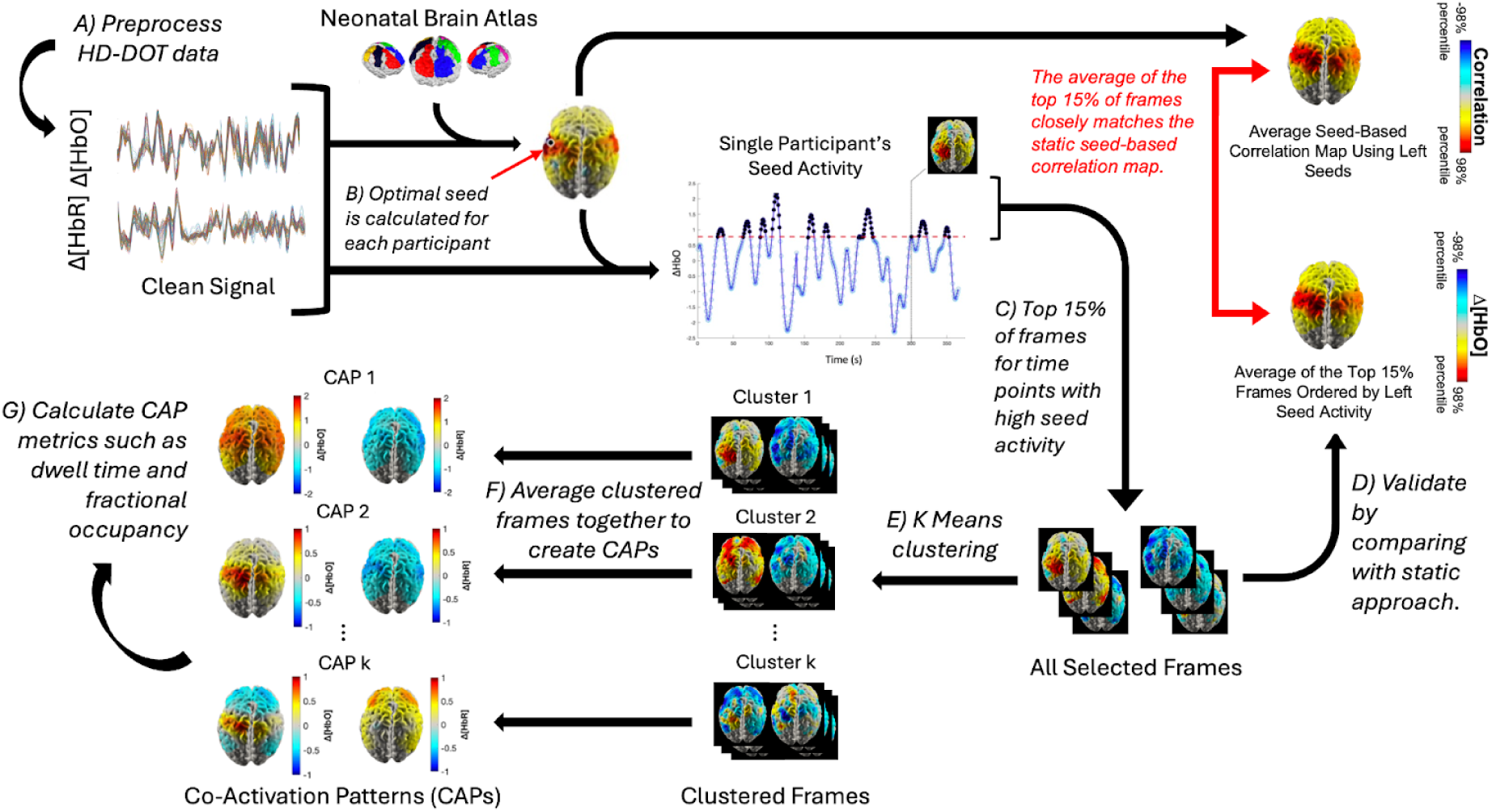
Analysis pipeline from preprocessing to CAP generation.

### 2.6 Frame Selection

To isolate time points that contribute most strongly to region-specific CAPs, data frames were selected based on seed activity (Figure 4). This procedure follows Liu and Duyn’s pre-clustering procedure for fMRI CAP analysis, which focuses on moments of pronounced seed activations to reveal transient network configurations (Liu and Duyn, 2013). Datasets were first z-score normalized within participants to reduce the influence of inter-participant variability. For each ROI, data frames were sorted by [HbO] activity at the left and right seed locations. The top 15% of frames were retained for each participant separately for the left and right seeds. This threshold was chosen because the average spatial map of the top 15% of frames was highly correlated with the corresponding seed-based correlation map (Figure 4. and Supplementary Figure 2), while including frames beyond this threshold provided little additional benefit. This threshold was also used to compare the dynamic CAP procedure in this dataset against a standard static approach (see Figure 3d). Since frontal, central, and parietal ROIs each had two seeds (left and right), the frame selection process was performed once using the left side seed and then again with the right seed. The left-seed-selected frames were combined with the right-seed-selected frames, so that each ROI had a single dataset containing all frames in which at least one of the two seeds showed high activity.

**Figure 4.**
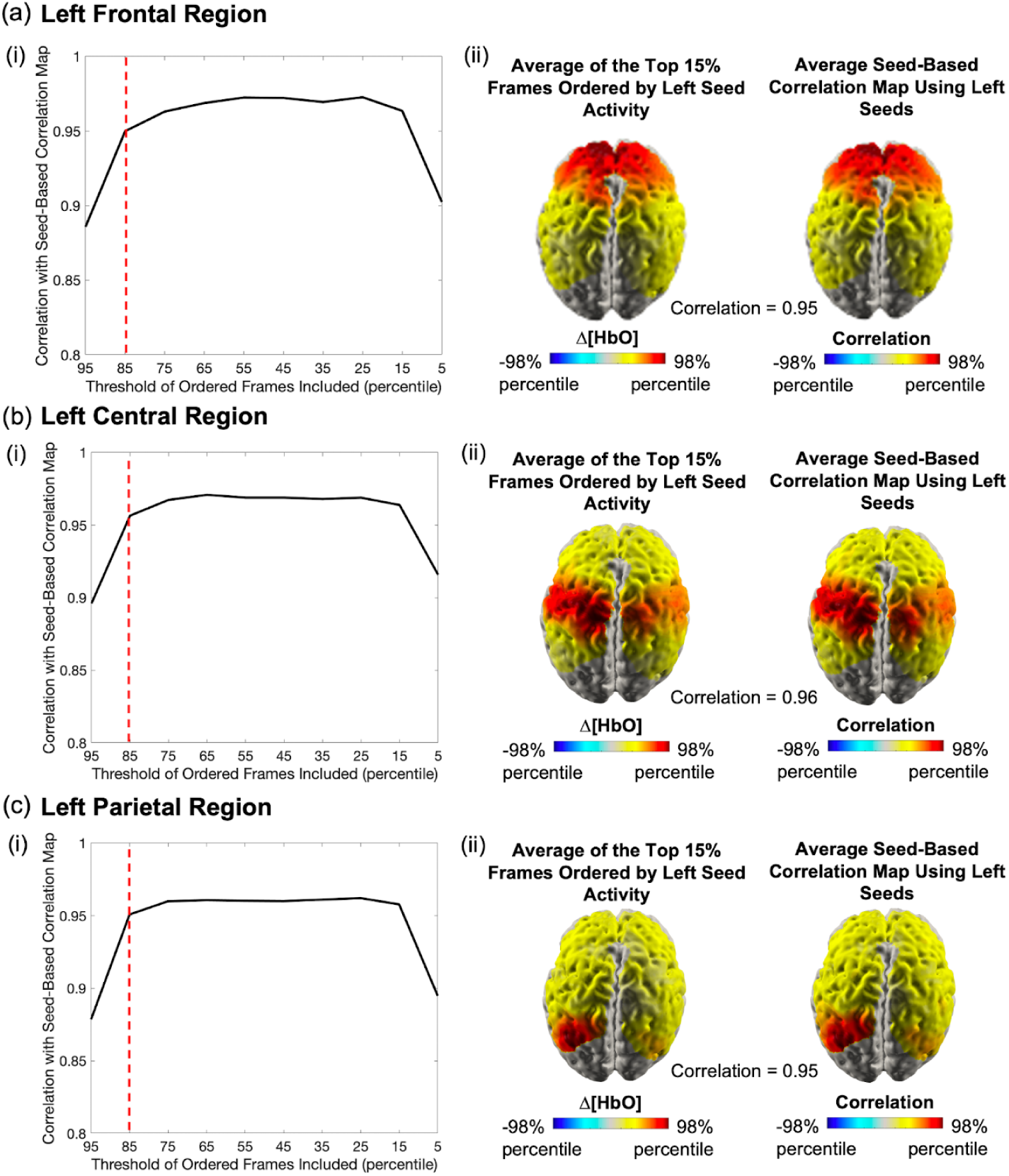
(i) Varying percentile threshold shows top 15% of frames ordered by seed activity is sufficient for high fidelity to static seed-based correlation analysis. (ii) Seed-based correlation maps are highly correlated with the spatial average of the top 15% of HD-DOT frames ordered by seed activity (correlation reported at the bottom of right-side figures). (a) Frontal left region seed analysis. (b) Central left region seed analysis. (c) Parietal left region seed analysis. Right equivalent region available in supplementary material (see Supplementary Figure 2)

### 2.7 K-Means Clustering

Since we wanted to take into consideration both [HbO] and [HbR] values from the 10984 sensitive nodes during clustering, we initially had a total of 21968 dimensions (10984 measures of [HbO] and 10984 measures of [HbR], see Figure 3e). Principal Component Analysis (PCA) is a popular dimensionality reduction algorithm used to improve the effectiveness and efficiency of unsupervised machine learning algorithms. PCA is an important preprocessing step for clustering problems with large datasets as it identifies linear combinations of the initial dimensions to reduce variables while retaining overall dataset variance. We used PCA from the Python module *sklearn.decomposition* to transform the group-level data along the node dimension from 21,968 nodes to 275 components, retaining 99% of total variance (Pedregosa et al., 2011). This transformation was done independently for each region’s dataset (frontal, central, and parietal).

PCA transformed datasets with dimensions of 275 × N, where N was the number of selected frames, were then clustered using the *KMeans* function from Python’s sklearn.cluster module (Pedregosa et al., 2011). K-means is an unsupervised machine learning algorithm which clusters data based on Euclidean distance. First, k centroids are randomly initialized. On each iteration of the algorithm, data points are assigned to their nearest centroid according to Euclidean distance. The centroid locations are then updated to reflect the mean of its associated data points. This step is repeated until the centroids remain stable between iterations (Pedregosa et al., 2011). Data points associated with the same centroid are said to be part of the same “cluster” of data. K-means is usually run several times since clustering results can be sensitive to the random initialization of the centroids. We repeated k-means analysis 100 times using the n_init parameter of sklearn’s *KMeans* function, which repeats k-means with different centroid seeds and returns the best run according to centroid inertia. Mean silhouette score, a metric commonly used to quantify cluster separation and similarity, was used to evaluate k-means performance. We tested a range of k (2–20) for each region independently and selected the most suitable k using the elbow method (see Supplementary Figure 3). Parameter k = 7 was selected for all three regions.

### 2.8 Co-Activation Patterns

Each cluster of similar frames corresponds to one CAP. The clustered frames were averaged together (in node space) to create the final CAP maps, which each have a Δ[HbO] representation and a corresponding Δ[HbR] representation (see Figure 3f).

We assessed CAPs by measuring the consistency, fractional occupancy, dwell time, and transition probability of each CAP (see Figure 3g) (França et al., 2024; Khan et al., 2022; Liu et al., 2018). CAP consistency refers to the intra-cluster spatial correlation of the CAP and is calculated by finding the mean spatial correlation of the CAP map with each frame in the cluster. Fractional occupancy, also known as occurrence rate or time fraction, is calculated at the participant level and refers to the proportion of frames in the CAP cluster to the total number of frames in the dataset. Dwell time is calculated at the participant level and refers to the mean duration of consecutive frames that belong to the CAP. Transition probability is determined at the participant level and refers to the likelihood of transitioning from CAP X to CAP Y. This is calculated by counting the number of transitions from CAP X to CAP Y and dividing it by the total number of transitions out of CAP X that occur in the dataset.

Though all four metrics characterize the CAPs, achieving high CAP consistency is precursory to further analysis because consistency quantifies how effectively the data were clustered and therefore how meaningful the CAPs are. Low consistency indicates poor data clustering and suggests CAP analysis is not suitable for the dataset. Using the CAP consistency values reported by Liu and Duyn (2013) as a benchmark (mean ± SD = 0.26 ± 0.04; median = 0.265; range = [0.21 – 0.30]), we found that CAPs in all regions achieved sufficient consistency for further analysis. In-participant fractional occupancy, dwell time, and transition probability were calculated for each CAP and averaged across participants.

## 3.0 Results

### 3.1 Frame Selection

For all regions, the average spatial map of the top 15% of frames ranked by seed activity showed a very high correlation (r ≥ 0.95) with the corresponding static seed-based correlation map (Figure 4; Supplementary Figure 2). This confirms that high-seed-activity frames provide an adequate representation of conventional seed-based functional connectivity and supports their use for subsequent CAP analysis.

### 3.2 Co-Activation Patterns

K-means analysis yielded clusters of varying sizes (see Supplementary Figure 4). A singular cluster from the central ROI analysis contained fewer than 50 frames, all sourced from the same participant and highly correlated. This cluster was excluded from further analysis because the small cluster size and single-participant origin suggested it was not a generalizable pattern. Frames within each cluster were averaged to generate the final CAP maps (see Figure 5 for z-scored [HbO] maps and Supplementary Figure 5 for complementary z-scored [HbR] maps). The CAPs are numbered according to their average consistency, from highest to lowest. Consistency scores for each region were as follows: Frontal CAPs: mean ± SD = 0.51 ± 0.20, median = 0.53, range = [0.24, 0.82]; Central CAPs: mean ± SD = 0.48 ± 0.22, median = 0.49, range = [0.19, 0.80]; Parietal CAPs: mean ± SD = 0.48 ± 0.21, median = 0.51, range = [0.25, 0.80]. Individual CAP consistency meets or exceeds the thresholds defined in the benchmark study (Liu and Duyn, 2013), except for central CAP 6. Although this CAP shows a lower consistency score (mean = 0.19), it is included to facilitate a complete and transparent report of our analysis. The high consistency across the CAPs strongly supports the validity of the identified patterns.

**Figure 5.**
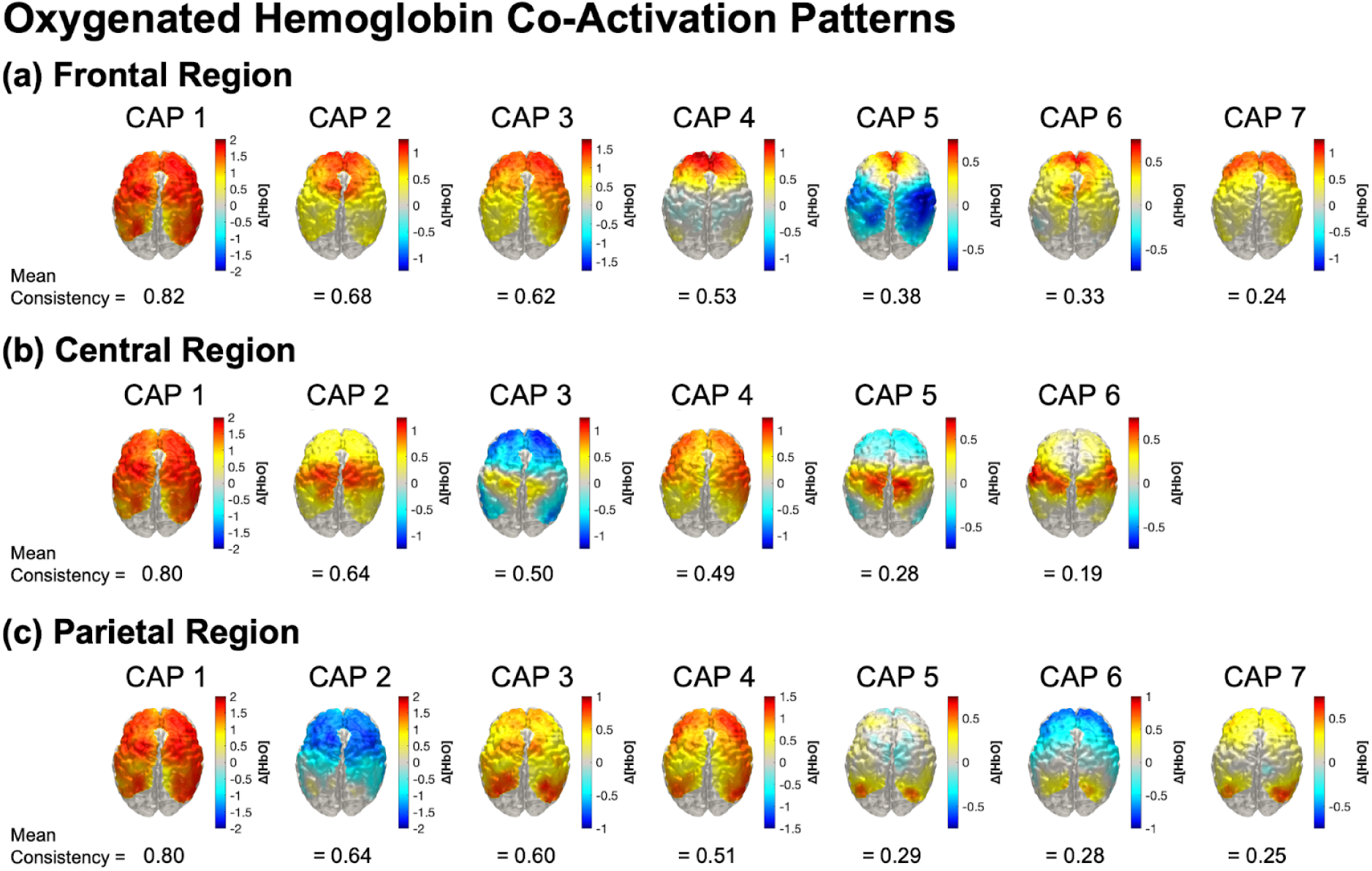
Z-score normalized oxygenated hemoglobin co-activation patterns (CAPs) for (a) frontal, (b) central, and (c) parietal regions. From left to right, the CAPs are shown in decreasing order of consistency. Each CAP’s deoxygenated hemoglobin counterpart map can be found in the supplementary material (Supplementary Figure 5)

To explore possible early formation of higher-order networks such as the DMN or FPNs, patterns exhibiting frontal and parietal co-activation were examined. Five CAPs from Figure 5 were visually identified to contain frontoparietal activity and are highlighted in Figure 6 across three views (frontal CAP 4, parietal CAP 3, parietal CAP 4, parietal CAP 5, and parietal CAP 7).

**Figure 6.**
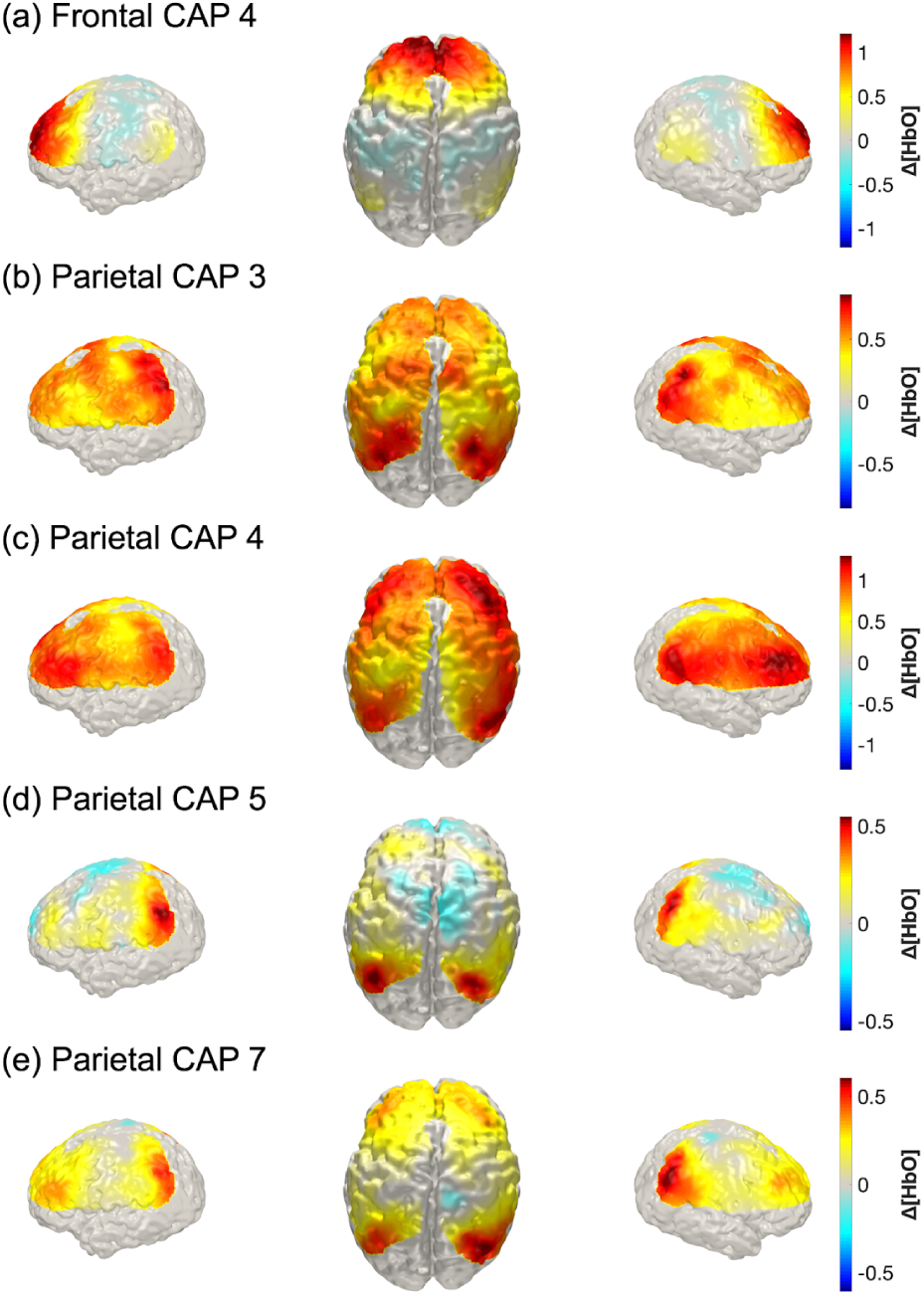
Three views (left lateral, dorsal, and right lateral) of z-score normalized Δ[HbO] CAPs with frontoparietal activity. Dorsal views for deoxygenated hemoglobin counterparts can be found in the supplementary material (Supplementary Figure 5).

### 3.3 Intra-Participant CAP Metrics

Metrics associated with the temporal features of each CAP, specifically fractional occupancy and dwell time, were explored in this study to check for significant differences across CAPs within each seed region (see Supplementary Table 1). All statistical tests were performed in R (R Core Team, 2017). A Shapiro-Wilk test was used to verify normality. Since normality was violated in all regions, a Friedman test was used to assess whether fractional occupancy and dwell time significantly differed between CAPs within each region while controlling for repeated measures within participants (see Supplementary Table 2). Fractional occupancy was found to be significantly different across CAPs for all three regions (p = 1.03e-05, p = 1.03e-15, p = 2.56e-23 for frontal, central, and parietal regions respectively), revealing that CAPs differ in occurrence rates. Dwell time was also found to be significantly different across CAPs for frontal and central regions (Bonferroni corrected p = 6.64e-04 and p = 5.91e-04 respectively) but not for the parietal region CAPs (Bonferroni corrected p = 0.203), revealing that CAPs persist for different durations. Pairwise differences were evaluated using a paired Wilcoxon test with Bonferroni correction (corrected for 21 comparisons for frontal and parietal regions and 15 comparisons for central regions). CAP metrics and significant differences with adjusted p values are shown in Figures 7 and 8.

**Figure 7.**
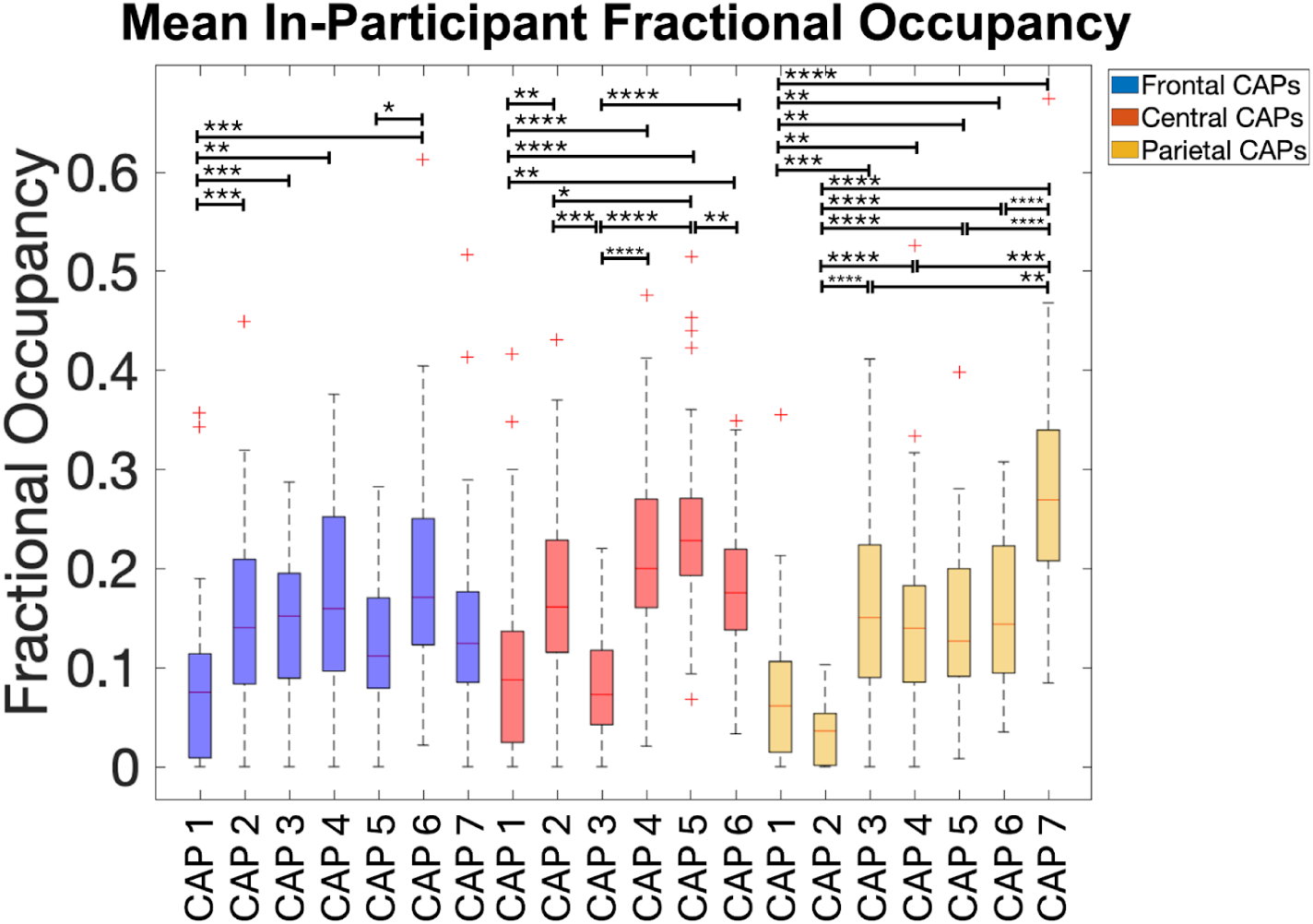
Average in-participant CAP fractional occupancies for all regions of interest. Bracketed pairs show significant statistical differences from Wilcoxon pairwise analysis. Bonferroni corrected: * p < 0.05; ** p < 0.01; *** p < 0.001; **** p < 0.0001.

**Figure 8.**
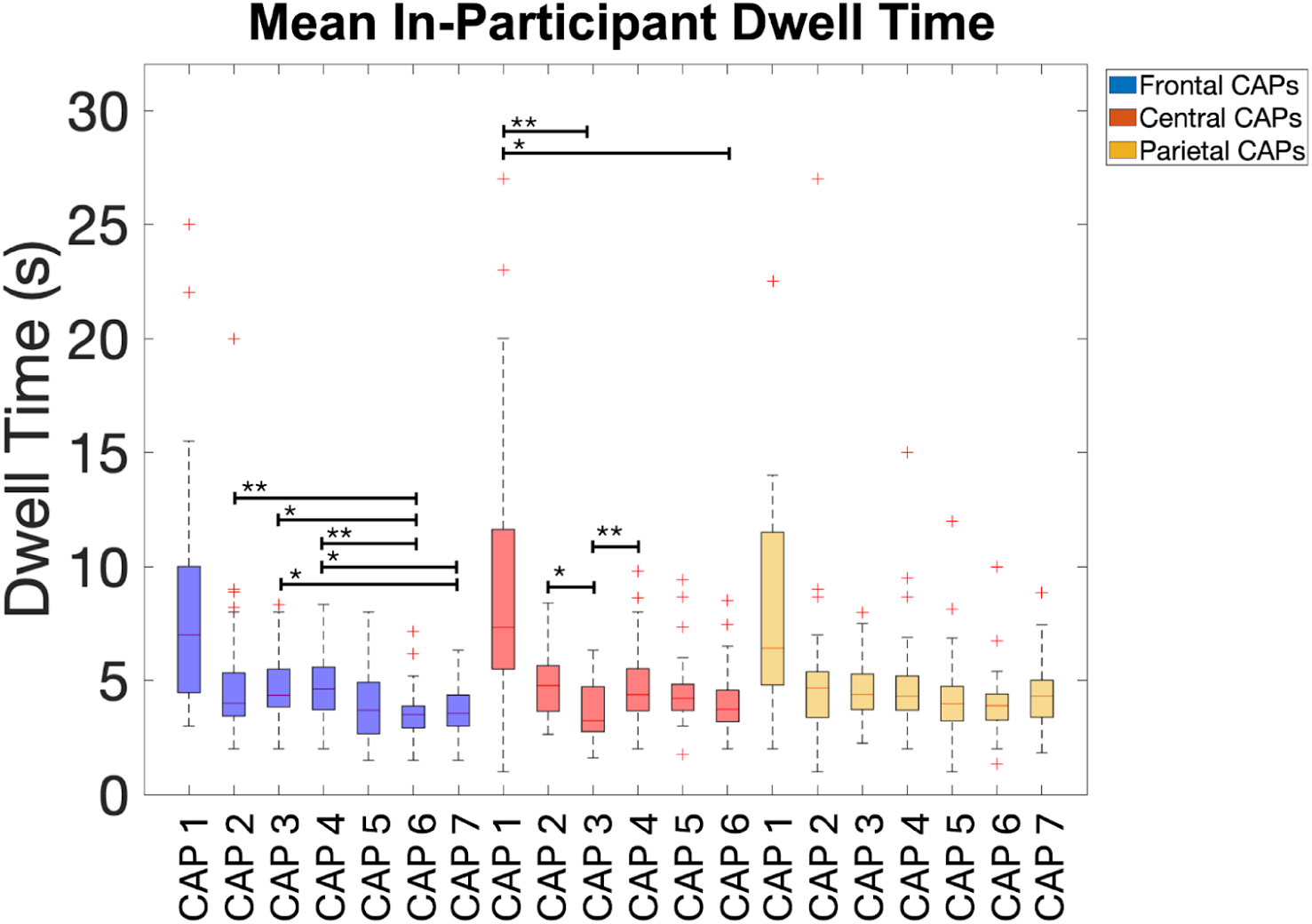
Average in-participant co-activation pattern (CAP) dwell times for all regions of interest. Bracketed pairs show significant statistical differences from Wilcoxon pairwise analysis. Bonferroni corrected: * p < 0.05; ** p < 0.01; *** p < 0.001; **** p < 0.0001.

Figure 9 presents the CAP transition probability matrices for frontal, central, and parietal regions. These matrices primarily demonstrate that the most probable transition is between frames of the same CAP, indicating most adjacent frames share membership. This was expected as fluctuations in concentration of HbO and HbR are slow, so most adjacent frames will have similar spatial patterns and therefore will likely be grouped into the same CAP cluster. Notably, these matrices are not symmetric along the diagonal.

**Figure 9.**
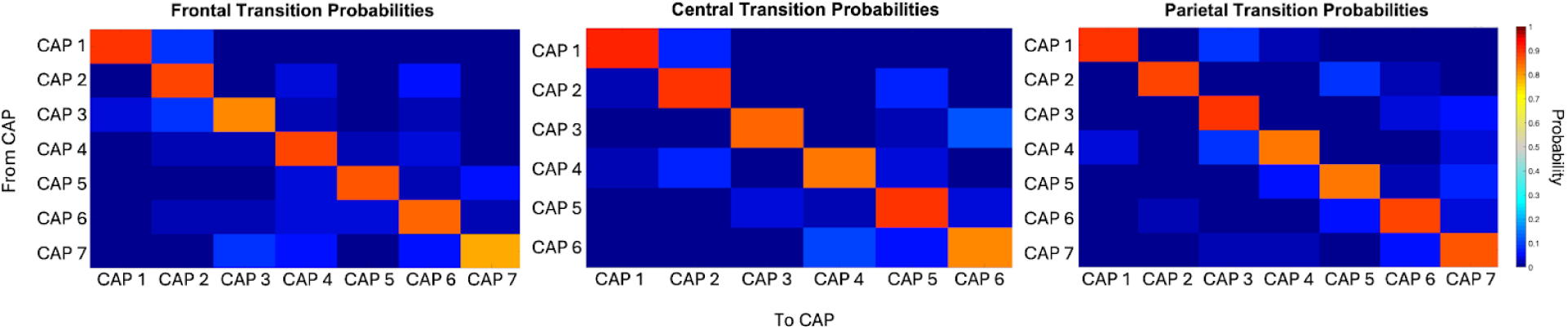
In-participant likelihood of transitioning to a given co-activation pattern (CAP) from each CAP for frontal, central, and parietal regions. Probabilities ≥ 5% are marked with a yellow circle.

## 4.0 Discussion

CAP analysis provides a time-resolved decomposition of activity across brain regions by identifying recurring spatial configurations expressed during high-activity moments. This decomposition is meant to complement and supplement our current understanding of functional connectivity which is largely derived from static analytical approaches. While static approaches average activity across an entire recording, CAPs separate the dataset into distinct recurring configurations that can differ in spatial arrangement, polarity (co-activation versus co-deactivation), and temporal expression. As a result, CAP analysis can demonstrate that an apparently single static connectivity pattern may in fact be composed of multiple transient states with distinct spatial and temporal features.

In this study, we applied CAP analysis to neonatal HD-DOT recordings which were acquired cot-side in sleeping term newborns. High activity frames from selected seeds closely resemble static seed-based connectivity maps, supporting the use of CAPs in this dataset. Clustering analysis revealed multiple recurring hemodynamic configurations within each region of interest, including seed-like patterns, anti-correlated configurations, and global increases in oxygenation. Notably, several frontal and parietal CAPs showed transient frontoparietal co-activation configurations consistent with early higher-order network organization that is not usually captured by static correlation maps.

### 4.1 Feasibility of CAPs in Neonatal HD-DOT

To the best of our knowledge, this is the first presentation of CAPs derived from neonatal brain activity, marking the first adaptation of the fMRI CAP analysis procedure proposed by Liu and Duyn for HD-DOT neonatal data (Liu and Duyn, 2013). Our primary goal was to establish the feasibility and demonstrate the suitability of neonatal HD-DOT data for CAP analysis. Before CAP clustering, we confirmed that frames with high seed activity were highly correlated with seed-based connectivity maps. This confirmation was critical, as CAP analysis relies on the assumption that top seed-selected frames are a sufficient proxy for seed connectivity. Verifying this assumption with HD-DOT data supports the feasibility of CAP analysis in this context and suggests it produces meaningful measures of functional connectivity.

The success of the clustering process itself, namely achieving mean consistency scores exceeding the fMRI benchmarks set by Liu and Duyn, further supports the compatibility of HD-DOT with CAP analysis (Liu and Duyn, 2013). While HD-DOT’s reduced spatial coverage compared to fMRI may partially explain these higher consistencies (fewer nodes reduce dimensionality and may improve clustering), the observed consistencies nonetheless indicate that neonatal HD-DOT data contain recurring, internally coherent spatial configurations that can be reliably identified using the CAP approach. Establishing the presence of these recurring patterns is significant because it suggests that traditional neonatal static seed correlation maps may be driven by repeating, transient co-activation configurations, consistent with adult fMRI CAP studies (Liu and Duyn, 2013).

### 4.2 CAP Interpretation

Although CAPs are identified by examining single-sample frames, they should be interpreted as recurring patterns of hemodynamic activity rather than instantaneous neural firing patterns. In each CAP, co-activating or deactivating areas represent a recurring relationship to the seeded region. This provides information beyond what is known by static FC methods because CAPs show the distinct spatial contexts in which a seeded region participates rather than only the average correlation pattern across the full recording.

The CAPs presented in this study primarily demonstrate symmetric activity across the left and right hemispheres. Visual inspection suggests three general trends across regional CAP sets. First, each set includes at least one CAP resembling the corresponding seed correlation map (e.g., frontal CAPs 2 and 7, central CAPs 2 and 6, parietal CAP 7). CAPs with similar overall patterns appear to differ mainly in magnitude and in which sub-regional areas are recruited (e.g., frontal CAP 7 appears relatively restricted to prefrontal activity, whereas frontal CAP 2 extends to dorsolateral prefrontal regions). Second, CAP sets typically include an anti-correlation pattern, showing positive Δ[HbO] in the seed region paired with negative Δ[HbO] outside the seed region (e.g., frontal CAP 5, central CAP 3, central CAP 5, and parietal CAP 6). Third, all CAP sets contain a pattern showing positive Δ[HbO] globally (e.g., frontal CAP 1, central CAP 1, and parietal CAP 1). This finding is consistent with other dynamic functional connectivity studies, including globally synchronous states identified in a neonatal fMRI study (França et al., 2024) and globally activated CAPs reported in a resting state rodent study (Matsui et al., 2016). Although non-neuronal physiological confounds such as respiration are attenuated during preprocessing, global hemodynamic fluctuations may still be present, and these CAPs may reflect instances of widespread increases in oxygenation across the imaged cortical surface.

### 4.3 CAP temporal characteristics and metrics

The metrics used to describe CAPs in this study (consistency, fractional occupancy, dwell time, and transition likelihood) were selected based on prior dynamic functional connectivity and related brain-state studies (Brown & Gartstein, 2023; França et al., 2024; Khan et al., 2022; Liu et al., 2018). It should be noted that hemodynamic coupling and downsampling to 1 Hz naturally promotes temporal smoothness, which may contribute to the multi-second dwell times and high self-transition probabilities observed in this study.

CAPs with global positive Δ[HbO] demonstrated the highest intra-cluster consistency within each set. This is potentially due to their relatively simple spatial organization, which may reduce variability across frames and participants. CAPs with lower consistency scores may reflect greater spatial heterogeneity within clusters and/or the presence of outlier frames that do not strongly resemble a dominant recurring configuration. Future work may benefit from formally identifying and excluding outlier frames prior to clustering to potentially improve within-cluster consistency. Fractional occupancy of the global positive Δ[HbO] CAPs appears significantly lower than several other CAPs within each region. CAPs with significantly lower fractional occupancies also tend to include the most negative Δ[HbO] patterns (frontal CAP 5, central CAP 3, and parietal CAP 2). Mean dwell time across CAPs was on the order of 3–5 seconds, and the global positive Δ[HbO] CAP frequently displayed a wider spread of higher dwell times, although this difference was only significant in the central region for two CAPs. Transition likelihood matrices suggest that frames most often transition reflexively, an expected finding given the timescale of hemodynamic dynamics. Although this study did not assess statistical significance of specific transitions due to the large number of comparisons and the absence of a priori hypotheses, visual inspection shows that some CAPs preferentially transition to other CAPs in a specific and directed way, suggesting that CAPs are not all equally related to one another. The transition matrices are also not symmetric along the diagonal, suggesting that when CAPs are weakly associated, the relationship may be directionally biased. These temporal metrics are a key advantage of CAP analysis when compared with static FC methods, as they not only identify the spatial configurations that exist, but also how often they occur, how long they persist, and how they transition between one another.

### 4.4 Relationship to known neonatal functional connectivity networks

These findings complement the existing literature on neonatal functional connectivity networks. The traditional static analysis (group-level seed correlation activity maps; Figure 4 and Supplementary Figure 2) closely resembles patterns reported in prior neonatal studies of static functional connectivity (Fransson et al., 2007; Gao et al., 2017; Keunen et al., 2017; Smyser et al., 2010). Comparison of CAP spatial arrangements with a functional parcellation of the neonatal cortical surface (Myers et al., 2024), as well as static neonatal functional connectivity studies, suggests similarities between CAPs and known neonatal networks. Specifically, frontal CAP activity resembles descriptions of salience network activity reported in one-month-old infants (Gao et al., 2017); central CAP activity aligns with descriptions of the sensorimotor network observed in early neonatal resting-state studies (Fransson et al., 2007); and parietal CAP activity is consistent with descriptions of modular and immature frontoparietal networks (FPNs) and default mode network (DMN) organization. Critically, CAP analysis shows that these network-like patterns occur intermittently in this neonatal population, rather than being expressed continuously as a time-averaged static functional connectivity map may imply.

### 4.5 Frontoparietal CAPs and higher-order network interpretations

The frontal and parietal CAPs are of particular interest because they may provide insight into the early emergence of higher-order networks such as the default mode network (DMN) and frontoparietal network (FPN), which can be difficult to characterize using static approaches. Prior neonatal fMRI studies have reported partial or immature DMN-like organization early in life, although the timing and extent of frontoparietal involvement remain variable across studies (Doria et al., 2010; Gao et al., 2017). In this study, we identified CAP configurations that transiently resemble medial frontal–lateral parietal co-activation. Given that medial frontal and lateral parietal connectivity is a key component of the mature DMN, these CAPs suggest features of later DMN organization may be expressed intermittently at term age, a finding not as readily apparent from static FC measures.

Several CAPs (notably frontal CAP 4 and parietal CAP 3) show co-activation between medial frontal and lateral parietal cortical regions (Figure 6a–b). In addition, some CAPs appear reminiscent of more modular DMN components: frontal CAPs 5 and 6 show relatively focused medial frontal activation (consistent with anterior DMN components), while parietal CAP 6 shows comparatively stronger posterior parietal activation (consistent with posterior DMN components).

Early in life, FPNs are immature and difficult to differentiate from DMN due to overlapping lateral parietal regions. One useful distinction is that DMN frontal involvement is more medial when compared with FPN involvement which is more lateral. CAPs with concurrent lateral frontal and lateral parietal involvement (parietal CAPs 4, 5, and 7; Figure 6c–e) may therefore reflect early FPN-like configurations. Parietal CAP 5 is particularly notable because the medial frontal region associated with DMN-like activity is deactivated, consistent with a configuration resembling FPN expression paired with DMN suppression.

Lastly, although parietal CAP 3 is discussed above as potentially reflecting DMN activity, its notable co-activation of central regions suggests possible overlap with attention-related networks. In mature adult brains, DMN and dorsal attention network (DAN) activity is typically anti-correlated. This anti-correlation, however, has been shown to decrease during sleep (Picchioni et al., 2013). Because neonates in this study were sleeping, transient co-activation of regions that later become part of opposing networks may reflect developmental immaturity, sleep-dependent organization, or both.

Overall, our dynamic functional connectivity analysis suggests that frontoparietal connections relevant to higher-order networks can be expressed transiently at term age. These results are consistent with neonatal fMRI studies reporting frontoparietal patterns using other dynamic approaches (França et al., 2024; Ma et al., 2020). The frontoparietal CAPs highlight transient configurations that are not emphasiszd by static correlation maps and may be especially informative for studying early higher-order network organization.

### 4.6 Addressing Challenges in Neonatal Imaging

CAP analysis paired with HD-DOT offers significant advantages for functional imaging brain studies in neonates and is a promising avenue for future dynamic functional connectivity studies. As previous studies (Collins-Jones et al., 2024; Frijia et al., 2021; Uchitel et al., 2023) have established, HD-DOT is particularly fit for imaging the infant cortex, as infant brains are smaller and their skulls are thinner than those of adults, allowing the NIR light to penetrate more deeply. These studies highlight that the LUMO system is comfortable, wearable, partially motion-tolerant, and non-invasive, qualities that can enable longer and more naturalistic studies of infants (Collins-Jones et al., 2024; Frijia et al., 2021; Uchitel et al., 2023). This study validates an analysis method, CAP analysis, which can complement HD-DOT’s strengths because it is also resilient to challenges in neonatal brain imaging analysis. Unlike other methods that rely on long, continuous datasets (e.g., correlation, ICA, dynamic ICA), CAP analysis is effective even when limited to shorter or discontinuous data segments. This is a significant advantage, as neonatal datasets are often hindered by motion artifacts and interruptions due to infant non-compliance. By focusing on singular high-activity frames, CAP analysis bypasses the typical dependency on long data windows, making it an ideal tool for studying the infant brain.

Though this study focused on healthy term infants, HD-DOT is also suitable for imaging sick or preterm infants in neonatal intensive care. Methods capable of studying these vulnerable populations are a crucial prerequisite to monitoring and improving their clinical outcomes.

### 4.7 Methodological Considerations

#### 4.7.1 Global Signal Regression

Global signal regression (GSR) was not used in the original analysis because GSR may potentially introduce spurious anti-correlations (Anderson et al., 2011; Murphy & Fox, 2017), although a similar approach was implemented by regressing out the mean signal from short distance channels. Previous fMRI CAP studies show mixed results on whether GSR has a constructive effect on the patterns that emerge (Liu and Duyn, 2013a, 2013b). To explore the effect of GSR on our data, analysis was repeated with the addition of GSR post preprocessing and before data normalization. The resulting CAPs are documented in the supplemental material (see Supplementary Figure 6). Notably, the global high activity CAPs are no longer present, and anti-correlation is prominent in all patterns. While this study does not focus on the GSR results, as short separation regression sufficiently attenuates physiological confounds and there is little consensus on the use of GSR, future studies may consider using GSR to investigate the impact of global hemodynamic fluctuations on CAP analysis.

#### 4.7.2 Selecting k

Though k selection is empirically chosen and data-driven in this study, it was apparent during analysis that silhouette score is not an ideal metric to use in k selection due to the nature of brain data. Silhouette score is not only a measure of intra-cluster similarity but also inter-cluster separation. Even after processing with PCA, it is unclear whether we can reasonably expect well-separated clusters in spatial brain data, as states may be highly similar spatially but differ functionally. Even though we arrive at a compelling optimal k using silhouette scores and the elbow method, the silhouette scores themselves appear to be low. Since consistency, a measure of intra-cluster similarity, is high, we hypothesize that the silhouette scores are low because of poor cluster separation. Liu addresses a similar concern in their paper (Liu et al., 2018), hypothesizing that activity patterns do not need to be distinct in order to be meaningful. We agree that our results are hierarchical groupings of frames rather than clearly disparate clusters. Nevertheless, k selection is non-trivial, as choosing a too high k increases redundancy of CAPs and choosing a too low k washes out potential nuances by combining clusters that should be distinct. Selecting a data-driven k optimizes these two opposing risks. We use CAP consistency to determine whether a pattern in the dataset is reliable. Thus, even if silhouette scores are low, sufficient consistency allows us to be confident in our clusters.

### 4.8 Limitations

#### 4.8.1 Recruiting

As with most neonatal studies, data collection was limited due to inherent difficulties recruiting and studying newborn infants. Studies were conducted opportunistically cot-side in between feeds, so data collection was sometimes interrupted by unavoidable circumstances such as the baby waking and needing consoling, the family having visitors, or the family discharging from the hospital. Additionally, some infants naturally startled or fidgeted more than others, resulting in data loss. Compared to fMRI and EEG, HD-DOT is more resistant to motion, but it is still sensitive to gross movements, resulting in data loss.

#### 4.8.2 HD-DOT cap coverage

This study targeted particular regions to test HD-DOT CAP analysis in neonates. The LUMO caps were designed to specifically target these areas. Future studies may consider using a HD-DOT montage with whole-head coverage to examine other regions and networks of interest, particularly the occipital region/visual network and temporal region/auditory network, which we were not able to assess in this study.

#### 4.8.3 Methodological Standardization

Given that HD-DOT CAP neonatal research is still in early development, the field lacks consensus for several steps in the preprocessing and postprocessing pipeline, including data cleaning, seed selection, and defining data quality thresholds. Though this work references previous HD-DOT infant studies for hyperparameters during preprocessing, a few novel decisions were made which would benefit from standardization in the future. One example is that this study pilots data-driven seed selection done at the participant level. This study aims to establish a strong case for CAP analysis with HD-DOT, and its success underscores the need for further methodological standardization to enhance reproducibility and comparability across studies.

## 5.0 Conclusion

This study offers a novel perspective on neonatal brain activity, marking the first application of CAP analysis in neonatal HD-DOT data. Our findings not only confirm the suitability of CAP analysis for neonatal HD-DOT data but also contribute valuable dynamic representations of known resting-state networks early in life. Future work involving larger longitudinal datasets may reveal how resting-state dynamic functional connectivity evolves over time. Through understanding the developmental trajectory of functional networks, we will enhance our understanding of typical and atypical brain development in early infancy.

## Code and Data Availability

The data that support the findings of this article are not publicly available due to privacy concerns. Scripts can be requested from the author at. DOT-HUB scripts are available on GitHub (DOT-HUB (https://github.com/DOT-HUB).

## Author Contributions

**Katharine Lee:** Conceptualization, Methodology, Data Collection, Visualization, Analysis, Writing (draft and editing). **Topun Austin:** Investigation, Supervision, Conceptualization, Methodology, Writing (editing). **Julie Uchitel**: Methodology, Data Collection, Writing (editing). **Cesar Caballero-Gaudes:** Conceptualization. **Liam Collins-Jones:** Methodology, Writing (editing). **Rob Cooper and Jem Hebden:** Conceptualization. **Andrea Edwards and Kelle Pammenter:** Data Collection. **Grace Kromm:** Data Collection, Writing (editing). **Borja Blanco:** Supervision, Conceptualization, Methodology, Writing (editing).

## Informed Consent Statement

The National Research Ethics Service East of England Committee approved this study (REC reference 15/LO/0358). Written informed consent was provided by parents for inclusion in the study as well as photography for academic distribution.

## Acknowledgments

This study was funded by Action Medical Research (GN2859). The NIHR Cambridge Biomedical Research Centre (BRC) is a partnership between Cambridge University Hospitals NHS Foundation Trust and the University of Cambridge, funded by the National Institute for Health Research (NIHR). TA is supported by the NIHR Cambridge BRC and the NIHR HealthTech Research Centre for Brain Injury. The views expressed are those of the author(s) and not necessarily those of the NIHR or the Department of Health and Social Care.

BB was supported by the Medical Research Council Programme Grant MR/T003057/1 and UKRI Future Leaders fellowship (grant MR/S018425/1). KL is supported by a Gates Cambridge Scholarship.

## Supplementary Materials

**Supplementary Figure 1.**
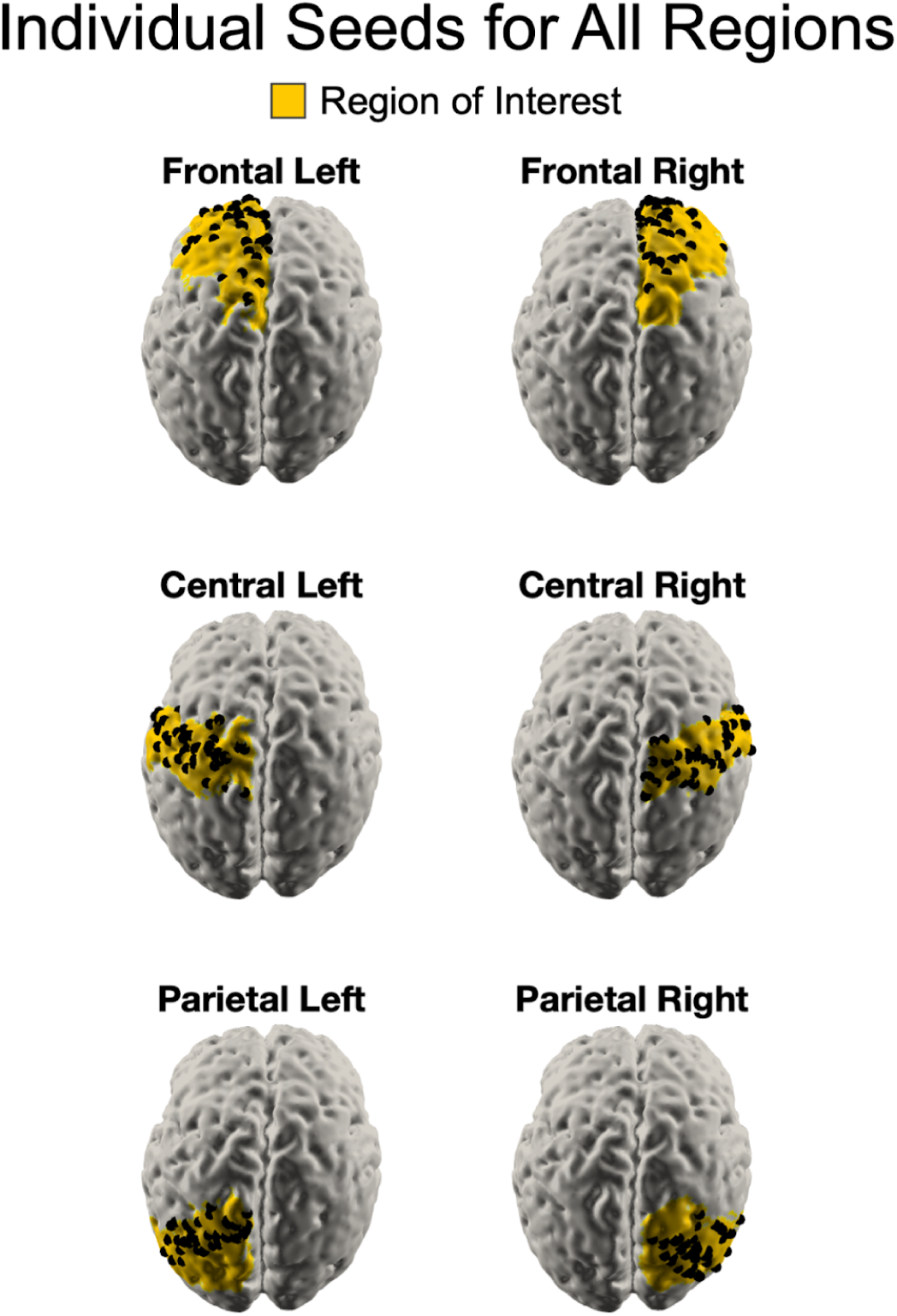
Optimal seed locations for each participant shown for all regions.

**Supplementary Figure 2.**
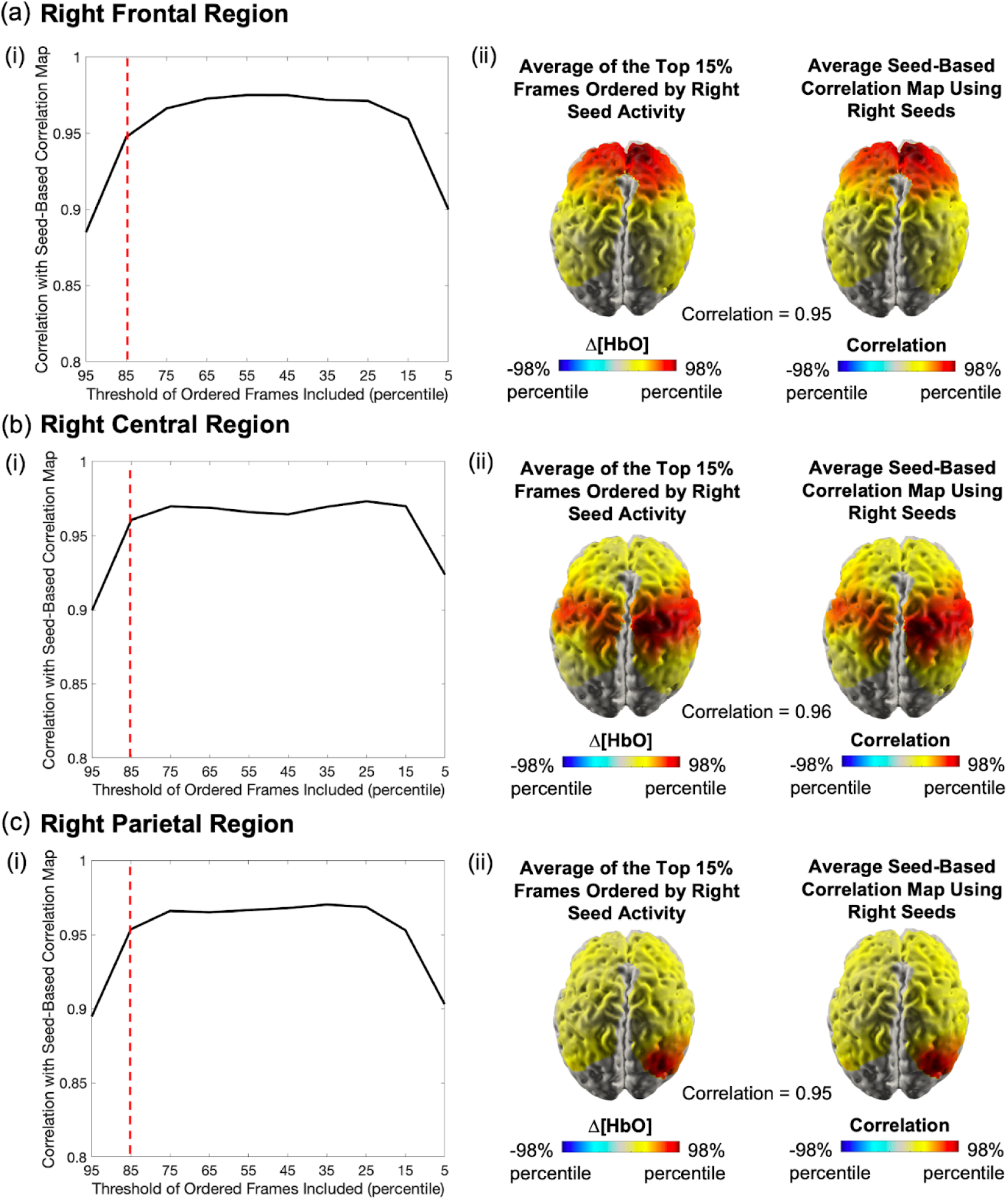
(i) Varying percentile threshold shows top 15% of frames ordered by seed activity is sufficient for high fidelity to static seed-based correlation analysis. (ii) Seed-based correlation maps are highly correlated with the spatial average of the top 15% of HD-DOT frames ordered by seed activity (correlation reported at the bottom of right figures). (a) Frontal right region seed analysis. (b) Central right region seed analysis. (c) Parietal right region seed analysis.

**Supplementary Figure 3.**
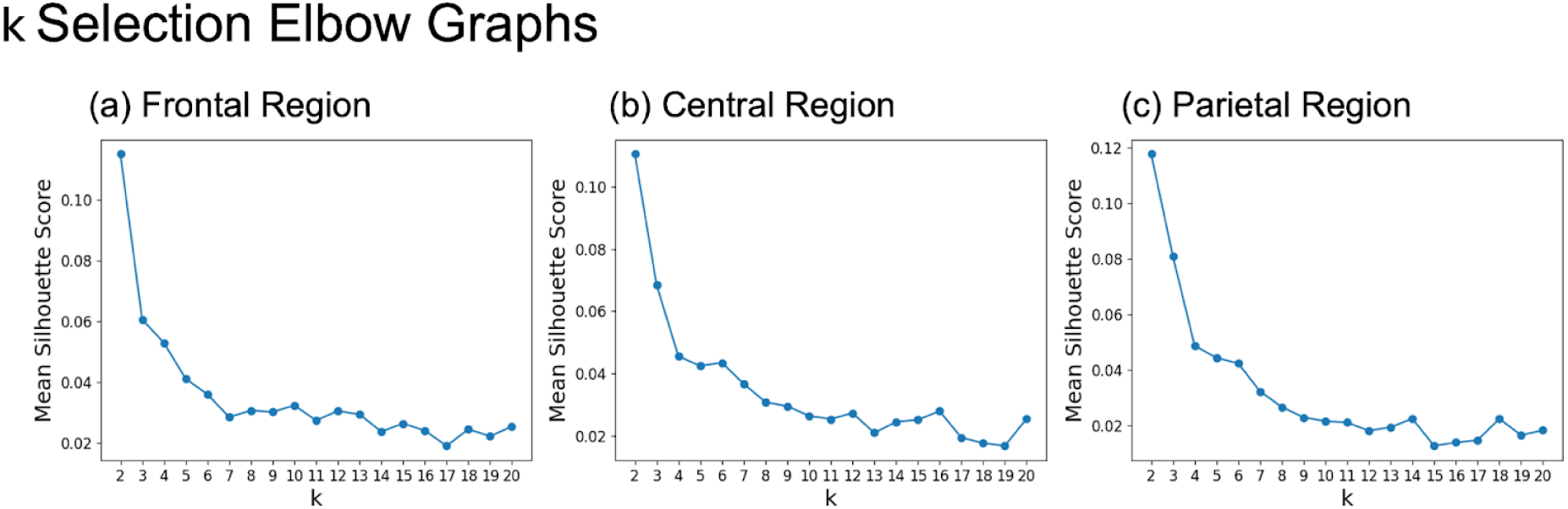
Elbow graphs depicting mean silhouette scores for each run of K-Means (k = [2, 20]) for (a) frontal, (b) central, and (c) parietal regions. Point of inflection was used to choose the best value for k.

**Supplementary Figure 4.**
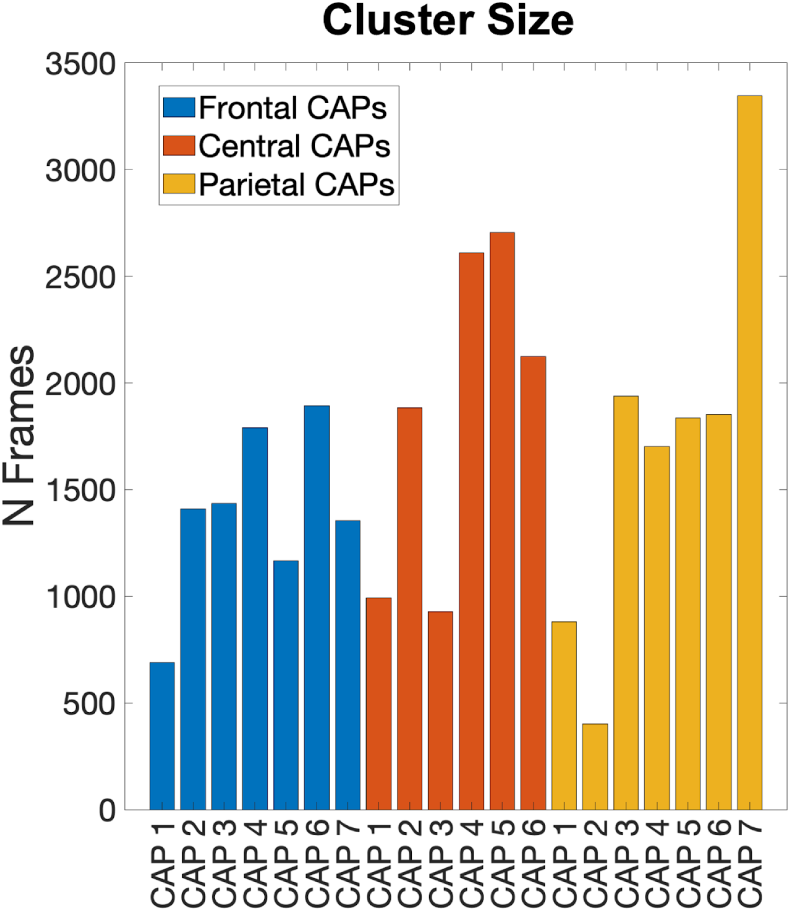
Number of frames in each CAP cluster.

**Supplementary Figure 5.**
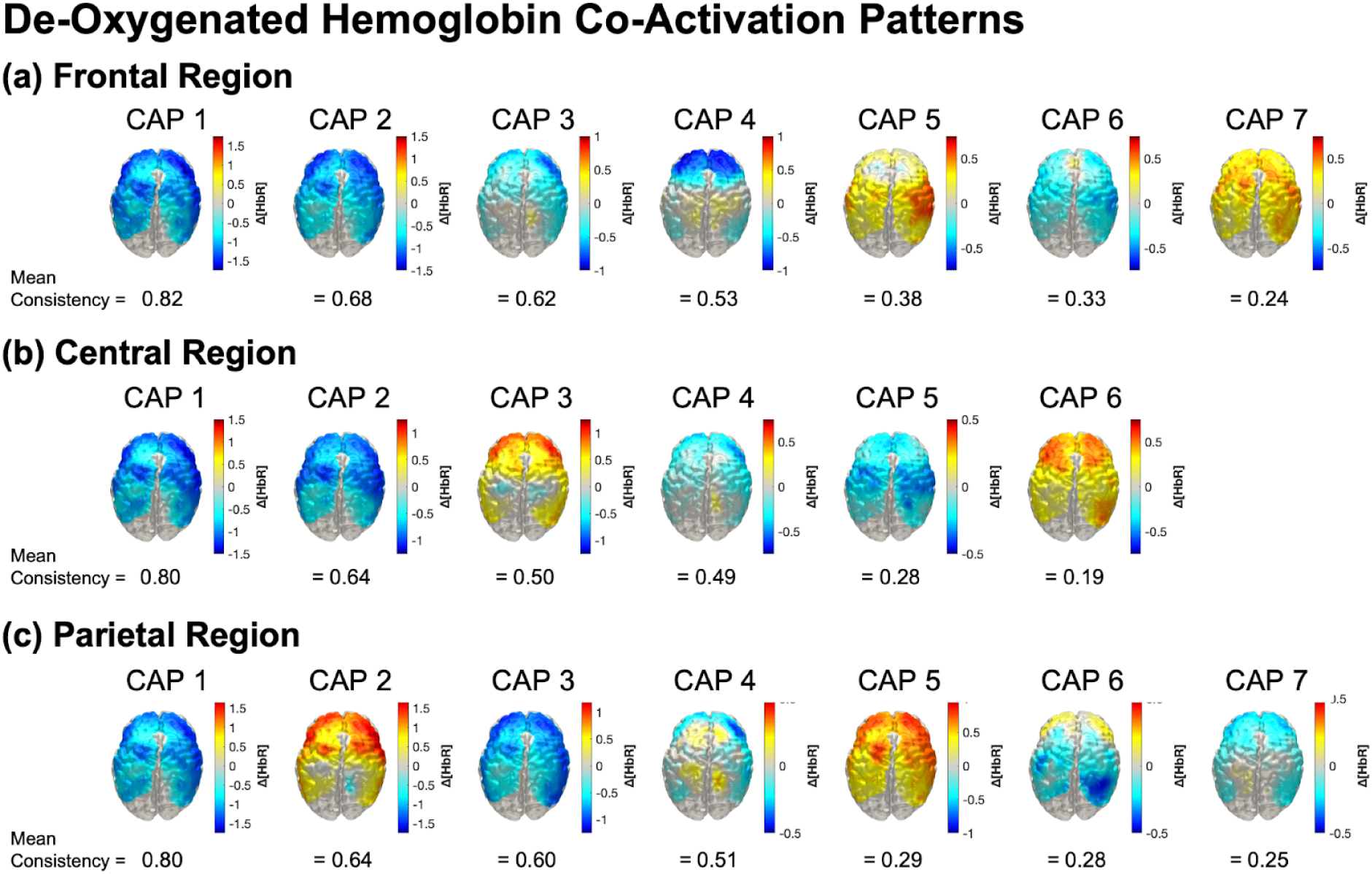
Z-score normalized deoxygenated hemoglobin CAPs for (a) frontal, (b) central, and (c) parietal region analyses. From left to right, the CAPs are shown in decreasing order of consistency.

**Supplementary Figure 6.**
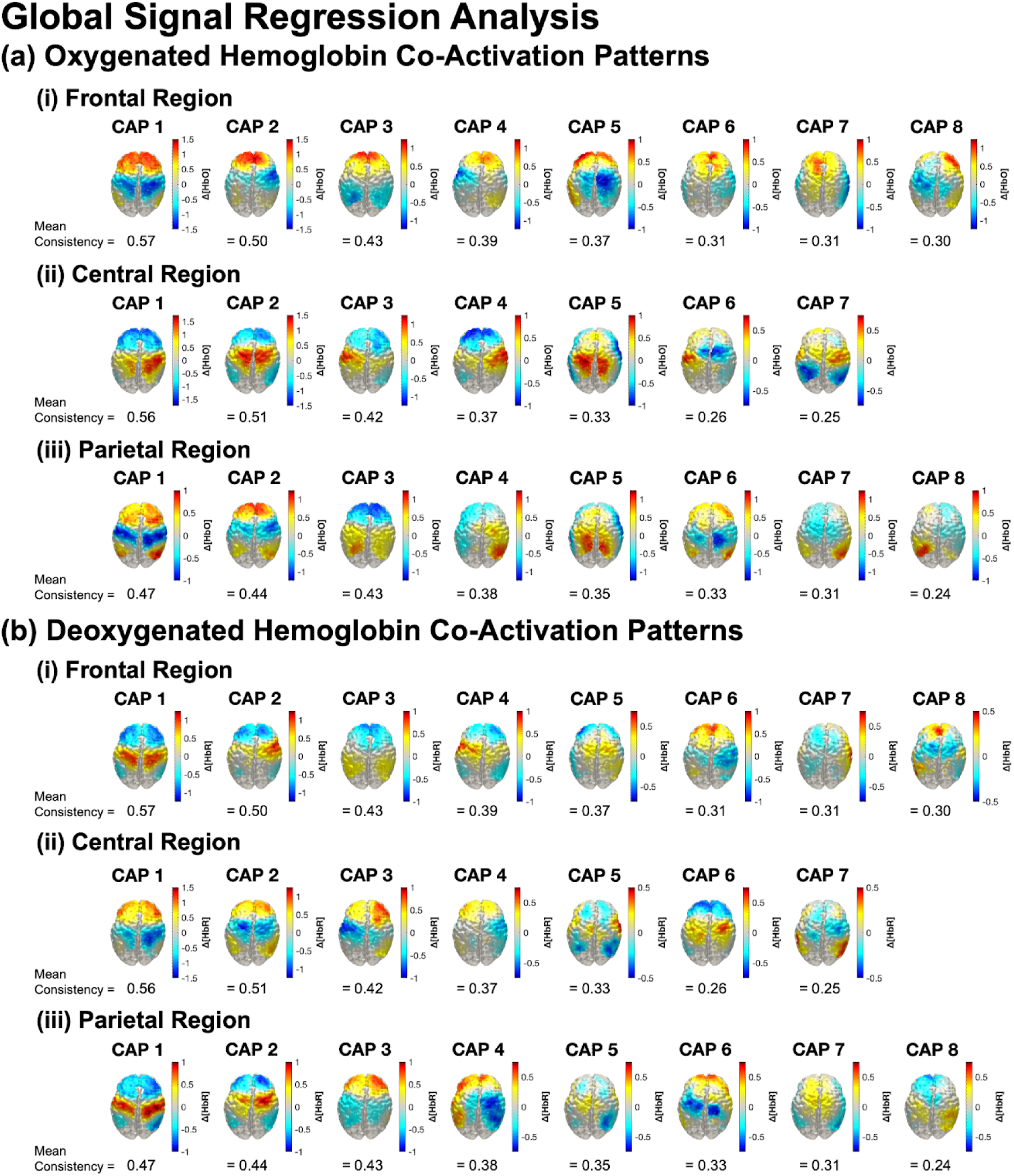
Results of analysis with global signal regression. (a) Z-score normalized oxygenated hemoglobin CAPs for (i) frontal, (ii) central, and (iii) parietal region analyses. (b) Z-score normalized deoxygenated hemoglobin CAPs for (i) frontal, (ii) central, and (iii) parietal region analyses. From left to right, the CAPs are shown in decreasing order of consistency.

**Supplementary Table 1.** CAP in-participant metrics (fractional occupancy and dwell time) for each region. Co-Activation Pattern (CAP) Metrics: Fractional Occupancy and Dwell Time Across Regions of Interest

| Region | CAP State | Fractional Occupancy (%) | Dwell Time (s) |
| --- | --- | --- | --- |
| Frontal | CAP 1 | 8.08 ± 8.31% | 8.21 ± 5.17 |
| | CAP 2 | 14.77 ± 9.10% | 4.84 ± 2.93 |
| | CAP 3 | 14.57 ± 6.93% | 4.74 ± 1.52 |
| | CAP 4 | 17.15 ± 9.93% | 4.82 ± 1.61 |
| | CAP 5 | 12.07 ± 6.40% | 3.77 ± 1.44 |
| | CAP 6 | 19.40 ± 11.23% | 3.52 ± 1.06 |
| | CAP 7 | 13.96 ± 9.48% | 3.67 ± 1.04 |
| Central | CAP 1 | 9.83 ± 9.65% | 9.27 ± 5.89 |
| | CAP 2 | 16.90 ± 9.34% | 4.77 ± 1.43 |
| | CAP 3 | 8.37 ± 5.72% | 3.76 ± 1.32 |
| | CAP 4 | 21.92 ± 8.84% | 4.72 ± 1.61 |
| | CAP 5 | 24.38 ± 9.17% | 4.46 ± 1.40 |
| | CAP 6 | 18.60 ± 7.52% | 4.03 ± 1.30 |
| Parietal | CAP 1 | 7.49 ± 7.57% | 7.81 ± 4.31 |
| | CAP 2 | 3.38 ± 2.86% | 5.13 ± 4.32 |
| | CAP 3 | 16.47 ± 9.79% | 4.62 ± 1.36 |
| | CAP 4 | 14.64 ± 9.67% | 4.76 ± 2.21 |
| | CAP 5 | 14.80 ± 7.58% | 4.25 ± 1.80 |
| | CAP 6 | 15.33 ± 7.64% | 4.02 ± 1.40 |
| | CAP 7 | 27.89 ± 11.29% | 4.38 ± 1.41 |

**Supplementary Table 2.** Statistical results for CAP metrics with Bonferroni correction. Statistical Results with Bonferroni Correction for Fractional Occupancy (FO) and Dwell Time Across CAPs

| Metric | Frontal | Central | Parietal |
| --- | --- | --- | --- |
| FO $p$-value | $1.03 \times 10^{-5}$ | $1.03 \times 10^{-15}$ | $2.56 \times 10^{-23}$ |
| Dwell Time $p$-value | $6.64 \times 10^{-4}$ | $5.91 \times 10^{-4}$ | 0.203 |

